# Evaluating the utility of amino acid similarity-aware kmers to represent TCR repertoires for classification

**DOI:** 10.1101/2024.12.06.626025

**Authors:** Hannah Kockelbergh, Shelley C. Evans, Liam Brierley, Peter L. Green, Andrea L. Jorgensen, Elizabeth J. Soilleux, Anna Fowler

## Abstract

Insights gained through interpretation of models trained on the T-cell receptor (TCR) repertoire contribute to advances in understanding of immune-mediated disease. This has the potential to improve diagnostic tests and treatments, particularly for autoimmune diseases. However, TCR repertoire datasets with samples from donors of known autoimmune disease status generally include orders of magnitude fewer samples than TCR sequences. Promising TCR repertoire classification approaches consider relationships between non-identical TCR sequences. In particular, kmer methods demonstrate strong and stable performance for small datasets. We propose a TCR repertoire representation that considers the relationships between amino acids within kmers flexibly and efficiently, which makes exploration of a wide range of TCR sequence features feasible. XGBoost models are trained and tested on kmer representations of TCR repertoire datasets including samples from patients with coeliac disease as well as donors with previous cytomegalovirus infection. We show that kmers that use small representative alphabets of amino acids are capable of training models that perform similarly or better than kmers based on all 20 amino acids. We find that, for cytomegalovirus infection status classification, defining amino acid relationships using BLOSUM62 can lead to a model with stronger performance as compared to an Atchley factor definition. Finally, we detail kmers or motifs which are important in each classification model and highlight the challenge of training truly interpretable TCR repertoire classification models which, if overcome, could lead to biomarker discovery for autoimmune diseases.

**Author summary:** TCR repertoire classification models can provide valuable understanding of autoimmune diseases if they can accurately infer autoimmune disease status and are biologically interpretable. Based on a kmer representation of the TCR repertoire, which has been shown to be most appropriate to train classification models on smaller datasets, we develop a computationally efficient method of grouping amino acid sequences to add knowledge to immune status classification model inputs, and consider its effect on interpretability. We find that most of the 4mer-based feature types we tested perform well in combination with an XGBoost model, where some benefit may be gained by applying a greatly-reduced alphabet of amino acids based on BLOSUM62 for cytomegalovirus serostatus classification. Our proposed reduced alphabet methodology is an alternative to kmer clustering which allows more efficient exploration of amino acid relationships and results in a more interpretable feature space.

## Introduction

T cells bind to specific antigens via their T-cell receptor (TCR). A naive T cell may clonally expand on encounter of an antigen to which its TCR specifically binds, resulting in a clonal population of T cells expressing the same TCR. TCRs may be formed of an *α* and *β* chain encoded by the TRA and TRB loci, or *γ* and *δ* chain encoded by the TRG and TRD loci; T cells expressing a TCR of each type, respectively, are called *αβ* T cells or *γδ* T cells, respectively. The diversity of TCRs expressed by an individual’s T cells, called their TCR repertoire, is typically vast; it has been estimated that a TCR repertoire of a single individual may include 100 million unique TCR *β* chains [1]. This diversity is underpinned by somatic recombination of Variable (V), Diversity (D) (only for *β* and *δ* chains) and Joining (J) gene segments that form the TCR genes, with addition of a Constant gene segment [2]. The complementarity-determining region 3 (CDR3) at the junction of joined V, D and J gene segments is the most variable region of TCR-encoding genes, and is generally considered to be the most important contributor to TCR-antigen binding specificity [3]. Next generation sequencing has enabled vast libraries of CDR3 sequences encoding TCRs from many T cells to be profiled in bulk.

Comparison of TCR repertoire samples from individuals with and without an immune status of interest can provide improved understanding of immune-mediated conditions. This approach has been applied to viral infection [4–25], which includes a study of TCR repertoires in the context of cytomegalovirus (CMV) infection by Emerson et al. [4]. CMV infection is highly prevalent and generally asymptomatic. Though CMV infection is important to study due to clinical relevance in cases of organ transplantation, pregnancy and immunodeficiency, many studies utilise the Emerson et al. dataset to evaluate immune status classification models [7, 8, 10], likely due to its large size and inclusion of 2 separate cohorts.

Autoimmune disease has also been studied using the TCR repertoire [13, 14, 26–38]. Coeliac disease (CeD) is an underdiagnosed autoimmune disease triggered by dietary gluten, where CD4+ *αβ* T cells bind to gluten-derived peptides presented by certain HLA-DQ molecules, resulting in damage to the small intestine [39]. Diagnosis of CeD may require a confirmatory duodenal biopsy, but valid findings rely on adequate consumption of gluten for a period beforehand. In addition, probability that two specialist pathologists disagree on the classification of a biopsy is estimated to be at least 20% [40], indicating that this test is somewhat subjective. *γδ* T cell populations in gut tissue of CeD patients are different compared to those who do not have CeD [41], suggesting relevance of *γδ* T cells in disease. Further understanding of CeD, as could be obtained through interpretation of a TCR repertoire classification model, may lead to new diagnostic tests.

While some “public” TCR sequences, which are TCR sequences that are observed in more than one individual, have been identified that are associated with immune status [4, 5, 22, 24, 27], consideration of shared attributes of TCRs within and between individuals may enable non-identical CDR3 sequences associated with a condition to be more readily identified, especially with smaller sample sizes where “public” TCRs are less likely to be observed. Strategies including deep learning [6, 8], TCR clustering [15, 42] and approaches that split TCR sequences into subsequences of length k, or kmers [7, 43–46] have been proposed which enable TCR similarity to be considered. In this work, kmers are targeted for further development due to their suitability as a classification basis for high dimensional TCR repertoire datasets with a small number of samples [7].

Within the CDR3 region of a TCR, it has been proposed that similar amino acids may lead to shared binding capabilities [43, 46]. It follows that incorporating amino acid relationships, defined by their physicochemical properties or likelihood of substitution for one-another, within a kmer representation may lead to recognition of non-identical kmers that share the same function within a TCR repertoire. However, if that is the case, the underlying relationships between amino acids that may lead to shared function are unknown. Previously, Atchley factors [47] have been utilised to encode kmers [43–46, 48, 49] in two related approaches, which both represent a kmer using a vector, where 5 Atchley factors are concatenated for each amino acid in the kmer. Models are then trained on encoded kmers [43–45, 49] or clusters of encoded kmers [46]. A study which benchmarks these 2 immune status classification approaches suggests that they have comparable performance to each other, but are both surpassed by a kmer-based gradient-boosted tree model that does not consider amino acid properties [7]. Though it is suggested that amino acid property information may improve the performance of their gradient-boosted tree model, it remains unclear to what extent the kmer representation or classification model of choice is responsible for differences in classification performance [7].

In kmeans clustering as undertaken by Thomas et al., the number of clusters must be specified in advance. 10 and 100 clusters are tried in their original publication [46], which each imply different amino acid similarity thresholds. Since training time would be expected to scale in proportion to the number of clusters [50], rigorously assessing a wide range of number of clusters, and therefore kmer similarity thresholds, would be extremely time-consuming. The study by Katayama and Kobayashi also highlights long training time of over a week for the method by Ostmeyer et al., despite their final model being limited to capturing linear relationships between the kmer representation and immune status [7]. An kmer-based representation was therefore developed which enables efficient assessment of multiple amino acid similarity thresholds.

This work explores the utility of considering amino acid properties in a kmer representation of TCR repertoire for classification. A novel representation of the TCR repertoire based on kmers defined with a reduced alphabet is proposed, in which similar amino acids are deemed to be equivalent. Our approach enables efficient and flexible exploration of TCR repertoire representations, allowing a range of amino acid relationships to be considered within an immune status classification model. This TCR representation method is benchmarked against the kmer clustering by Thomas et al. and unaltered kmers with no encoding applied. Additional results are also presented with amino acid similarity defined using BLOSUM62 [51], which to our knowledge is the first use of BLOSUM in a kmer representation of the TCR repertoire. These representations are used to train XGBoost models for CMV infection status and CeD status which are each evaluated using separate testing datasets. Methodology is otherwise kept consistent for fair comparison. As a result this work provides evidence of the effect of amino acid similarity information on the utility of a kmer representation of the TCR repertoire, which may guide future methodological development for TCR repertoire representations for classification.

## Results

### A flexible definition of amino acid reduced alphabets

Atchley factors and BLOSUM62 are used to define relationships between the 20 naturally-occurring amino acids. Reduced alphabets are defined by considering amino acids equivalent if they share similarity, where amino acid similarity is determined by similarity of Atchley factors or substitution likelihood by BLOSUM62. In practice, any user-defined amino acid properties or substitution matrix could be employed. Hierarchical clustering of amino acids results in multiple solutions which are defined at different similarity thresholds, which allows size of the reduced alphabet to be tuned as a hyperparameter. When applied to a kmer representation of the TCR repertoire, this flexibly-defined reduced alphabet enables optimal amino acid relationships to be chosen for a TCR repertoire classification model.

For each of Atchley factors and BLOSUM62, reduced alphabets over a full range of similarity thresholds, and therefore of size 1 to 20, are shown in fig 1. For Atchley factors, 20 different clustering solutions exist over a range of distances. For BLOSUM62, 19 different clustering solutions exist due to immediate transition from 15 to 17 clusters at distance of 0.333: equivalent pairwise similarity is calculated between glutamic acid (E) and glutamine (Q), as well as lysine (K) and arginine (R). Each of these reduced alphabets is applied to a 4mer representation of each dataset.

**Fig 1.**
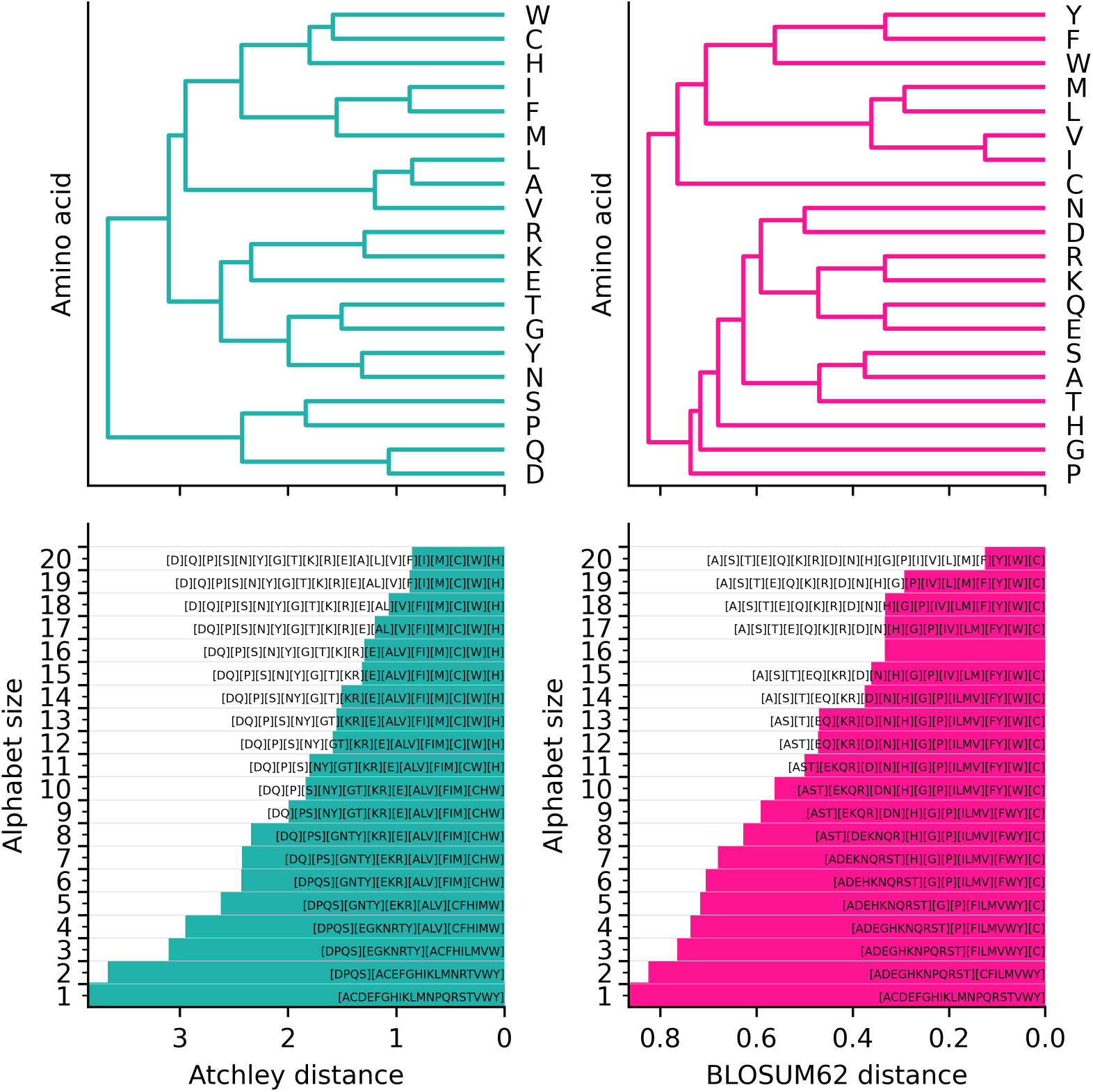
Reduced alphabets of amino acids. Atchley factor (left) and BLOSUM62-based (right) hierarchical clustering solutions for the 20 amino acids. Dendrogram top left and right show clusters of amino acids over a range of distance thresholds. Below, alphabet size is plotted over the distance threshold amino acid equivalence annotations. Amino acids considered equivalent within each alphabet are grouped using [].

In contrast, kmer clusters defined similarly as by Thomas et al. [46] define groups of whole kmers rather than amino acids. As in the original method, we use Atchley factors to calculate kmer clusters, and additionally BLOSUM62, with 100 clusters in each case. This fixed number of 100 clusters is used here since it found to result in a more sensitive and specific TCR repertoire classification model than with 10 clusters in the original publication [46]. Also, fitting the number of clusters as a hyperparameter over a full range of possible values is expected to lead to an insurmountable computational challenge due to the average computational complexity of kmeans clustering scaling linearly with the number of clusters [50]. If we assume that kmer clustering results in equivalent kmer groups as a reduced alphabet solution, the similarity threshold where amino acids would be considered equivalent corresponds to an alphabet size of 3.16. A standard kmer representation, similar to that by Katayama and Kobayashi [7] but with 4mers instead of 3mers and no end characters, corresponds to 4mers defined with an amino alphabet alphabet size of 20.

### CMV classification problem

The TCR *β* chain repertoire samples obtained from blood from 2 cohorts published by Emerson et al. were used as CMV training and testing datasets [4]. All 640 and 119 samples each within the training and testing datasets were downsampled to 23024 reads to avoid confounding by sample depth, with no loss of samples as a result. The CMV training and testing datasets are summarised in table 1.

**Table 1.** Summary of CMV datasets.

|  | CMV Training Dataset | CMV Testing Dataset |
| --- | --- | --- |
| Locus | TRB | TRB |
| Starting material | DNA from blood | DNA from blood |
| Samples before downsampling | 640 | 119 |
| CMV+ samples before downsampling | 289 | 51 |
| CMV- samples before downsampling | 351 | 68 |
| Downsampled reads to | 23024 | 23024 |
| Samples after downsampling | 640 | 119 |
| CMV+ samples after downsampling | 289 | 51 |
| CMV- samples after downsampling | 351 | 61 |

XGBoost models trained on 5 different kmer-based representations of the CMV training dataset are evaluated in validation, shown in table 2 and testing shown in table 3. For each evaluation, 6 performance measures are shown: area under receiver operating characteristic curve (AUROC), accuracy, sensitivity, specificity, balanced accuracy and Matthew’s correlation coefficient (MCC). In validation, the mean and standard deviation of each performance measure is calculated. Accuracy, though shown for completeness, is not a suitable measure on which to base our conclusions due to class imbalance in the CMV datasets. Each of sensitivity and specificity alone cannot fully capture model performance, but AUROC, balanced accuracy and MCC combine contributions from both of these measures into a single value. AUROC, shown in the first column of each table, is ultimately used to evaluate overall model performance for consistency with other work [7], and because it takes classification probability into account which is not the case for balanced accuracy or MCC.

**Table 2.**
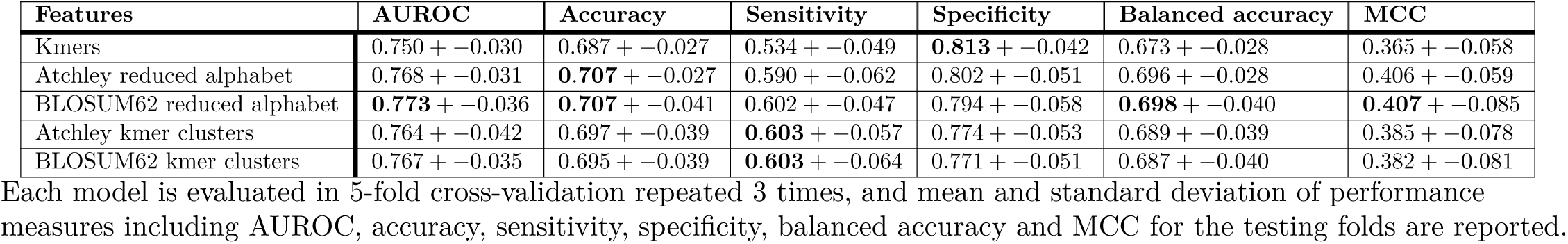
Validation performance of XGBoost models on CMV training datasets with five kmer representations.

**Table 3.** Testing performance of XGBoost models on CMV testing dataset with five kmer representations.

| Features | AUROC | Accuracy | Sensitivity | Specificity | Balanced accuracy | MCC |
| --- | --- | --- | --- | --- | --- | --- |
| Kmers | 0.768 | 0.681 | 0.706 | 0.662 | 0.684 | 0.364 |
| Atchley reduced alphabet | 0.781 | **0.731** | 0.667 | **0.779** | **0.723** | **0.449** |
| BLOSUM62 reduced alphabet | **0.802** | 0.697 | 0.647 | 0.735 | 0.691 | 0.382 |
| Atchley kmer clusters | 0.698 | 0.613 | 0.706 | 0.544 | 0.625 | 0.249 |
| BLOSUM62 kmer clusters | 0.760 | 0.664 | **0.765** | 0.588 | 0.676 | 0.352 |

Each model is trained on entire CMV training dataset and tested on entire CMV testing dataset. Performance measures including AUROC, accuracy, sensitivity, specificity, balanced accuracy and MCC are reported for testing dataset.

In validation, estimates of AUROC are similar across kmer feature types for an XGBoost model in validation on the CMV training dataset. A BLOSUM62 reduced alphabet results in the highest performance estimates across all measures reported except sensitivity and specificity, though an Atchley reduced alphabet results in performance that is very close to or equivalent, often with smaller error estimates. In addition, the standard kmer model AUROC is within 1 standard deviation of the estimated best-performing BLOSUM62 reduced alphabet model.

When all models were trained on the CMV training dataset and tested on the CMV testing dataset, a BLOSUM62 reduced alphabet is estimated to give the highest AUROC of 0.802, but is not universally best in terms of all measures. The model trained on Atchley reduced alphabet kmers also performs well, resulting in the highest balanced accuracy of 0.723 and highest MCC of 0.449. No one feature type gives rise to obviously superior performance in this classification task, though standard kmers and kmer clusters are judged to perform worse than reduced alphabets since they lead to lower AUROC, balanced accuracy and MCC. Despite lack of tuning of amino acid similarity threshold, kmer clusters can be used to train CMV classification models with an AUROC near 0.7 or above.

For the reduced alphabet kmer models, alphabet size is set as a hyperparameter such that AUROC is optimised. In fig 2, alphabet sizes between 5 and 14 are chosen across cross-validation iterations for both amino acid similarity types. Alphabet sizes of 10 are chosen within models that are trained on the entire CMV training set and evaluated in testing for Atchley and BLOSUM62 reduced alphabets. An alphabet size of 10 would be equivalent to 10^4^ = 10000 4mer groups assuming that all 4mers are observed, which is 2 orders of magnitude larger than the actual number of clusters used in kmer clustering.

**Fig 2.**
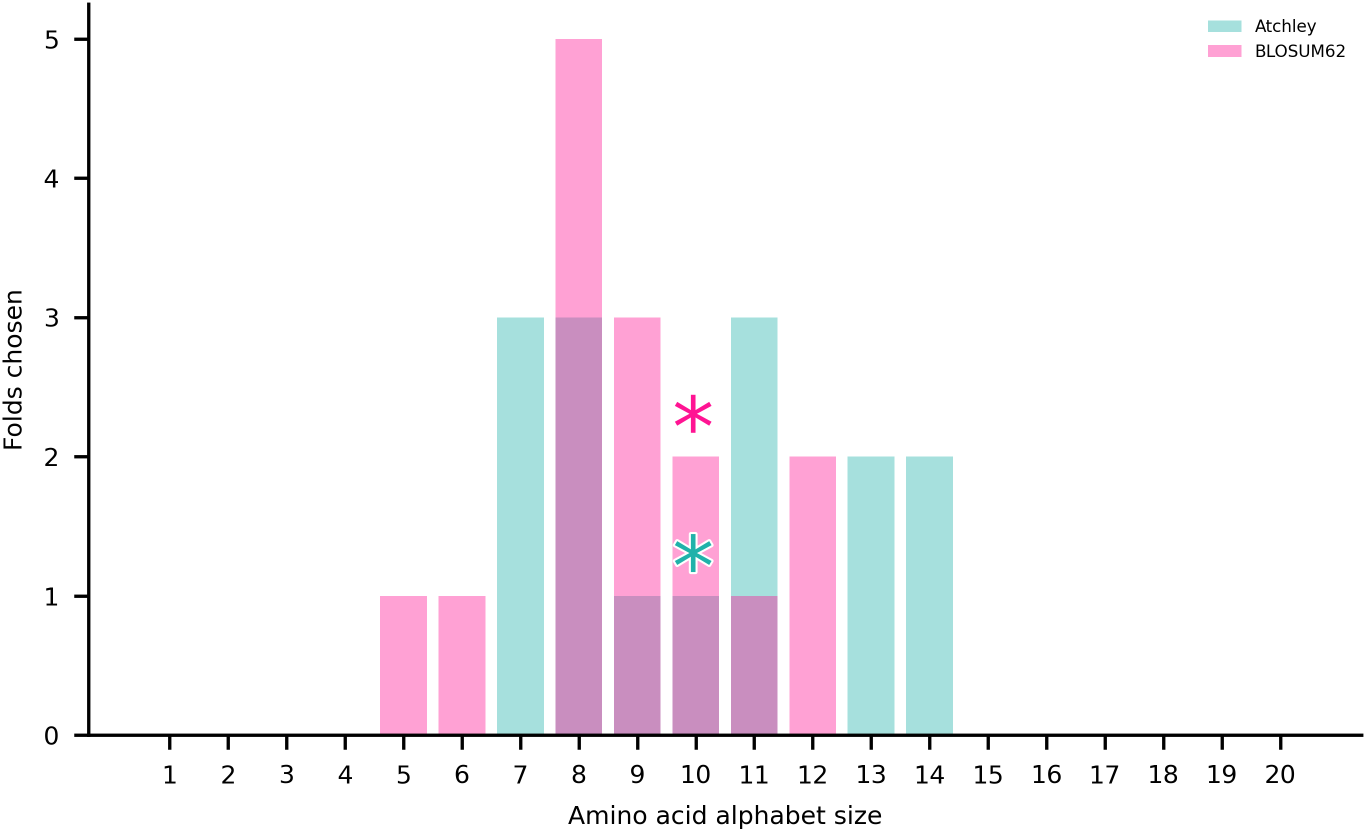
Reduced alphabet sizes chosen by XGBoost models in validation and testing with CeD datasets. Alphabet sizes chosen in training are indicated by asterisks.

We utilise the built-in feature importance method within XGBoost, specifically the gain, to calculate the contribution of each kmer-derived feature to the final models trained on the CMV training set. Where corresponding gain is zero, a feature is not utilised within an XGBoost model, and not included in these results. We also consider the number of kmers aggregated into each feature, and present kmer groups that result from application of a reduced alphabet; these groups are referred to as motifs. Fig 3 shows a summary of features including total number utilised by the model, proportion of importance, and number of kmers included within each feature for the XGBoost model trained on the entire CMV training set. For standard kmer and reduced alphabet kmer features, around 1000 or more features are incorporated into the model. In these cases, few features have a share of more than 1% of the total feature importance. Kmer cluster features tend to contain large numbers of kmers, and fewer are utilised within a model. A more detailed summary of features is included in S1 Fig, where kmers and kmer motifs that define each feature are annotated where possible. As shown in S1 Fig, a stacked bar plot showing feature importance as a proportion of the total, the motifs with the greatest feature importance within the reduced alphabet kmer models do not bear obvious similarity to the most important kmers within the standard kmer model. Since kmer clusters group together a large number of kmers, these features are described only with a numerical index for practical purposes. As such, comparison of the kmers included within these features to those in other feature types is not attempted.

**Fig 3.**
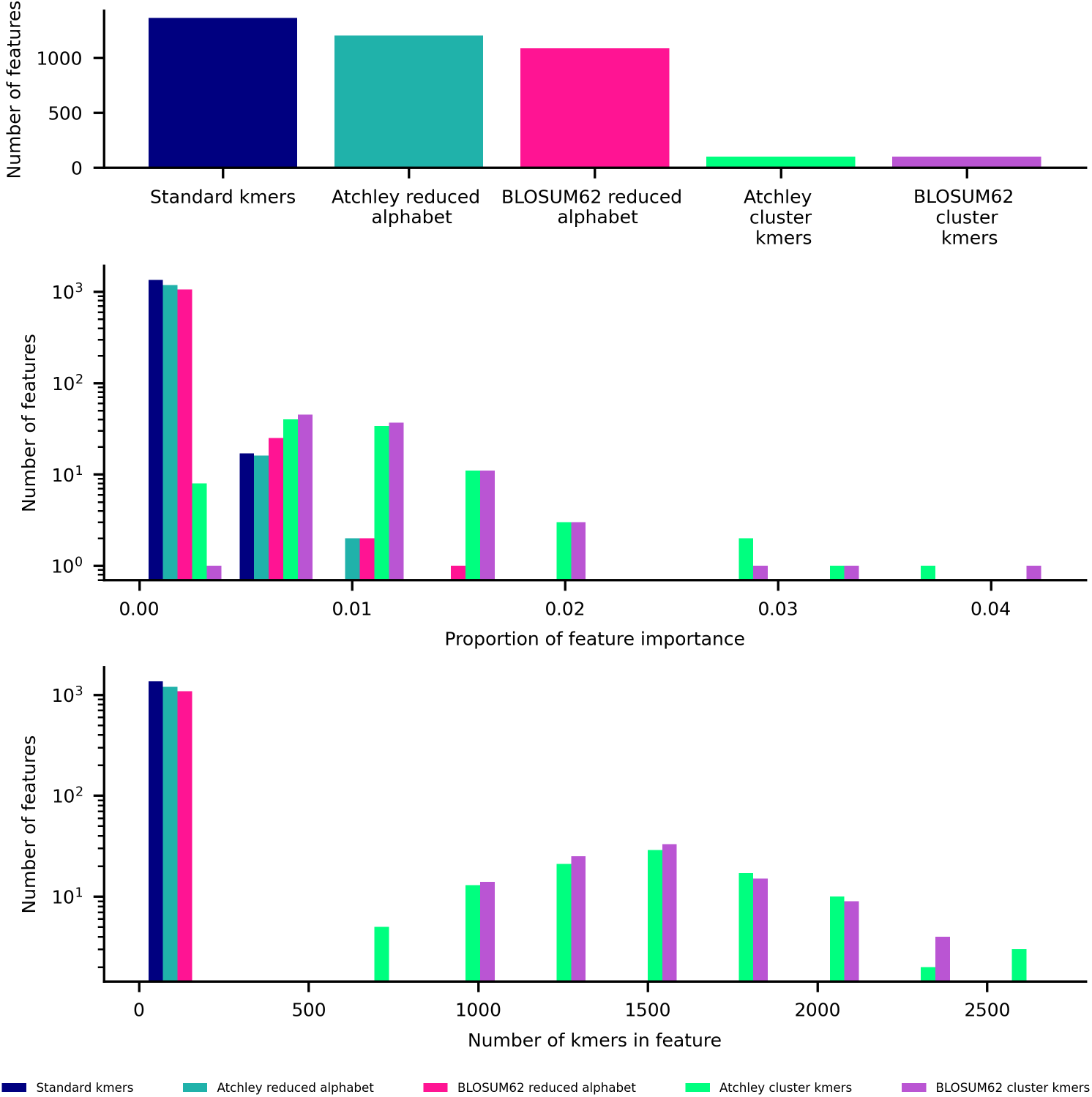
Summary of features used in XGBoost model trained on CMV training dataset for each feature type. Top: bar plot of number of features used in model. Middle: histogram of proportion of importance of features used in model. Lower: histogram of number of kmers within each feature used in model.

### CeD classification problem

A CeD training dataset which includes 139 *δ* chain TCR repertoire samples was obtained from duodenal biopsies. Another published *δ* chain dataset from duodenal biopsies was used as a CeD testing dataset [28]. Some samples with low TCR read counts in the CeD training dataset a possible source of confounding. A downsampling threshold was therefore chosen by trading off the number of samples retained with the read counts of each downsampled sample, illustrated by fig 4. The chosen downsampling threshold of 10078 was applied to both CeD datasets, resulting in a final count of 125 and 22 samples in each of the training and testing datasets respectively. The CeD training and testing datasets are summarised in table 4.

**Fig 4.**
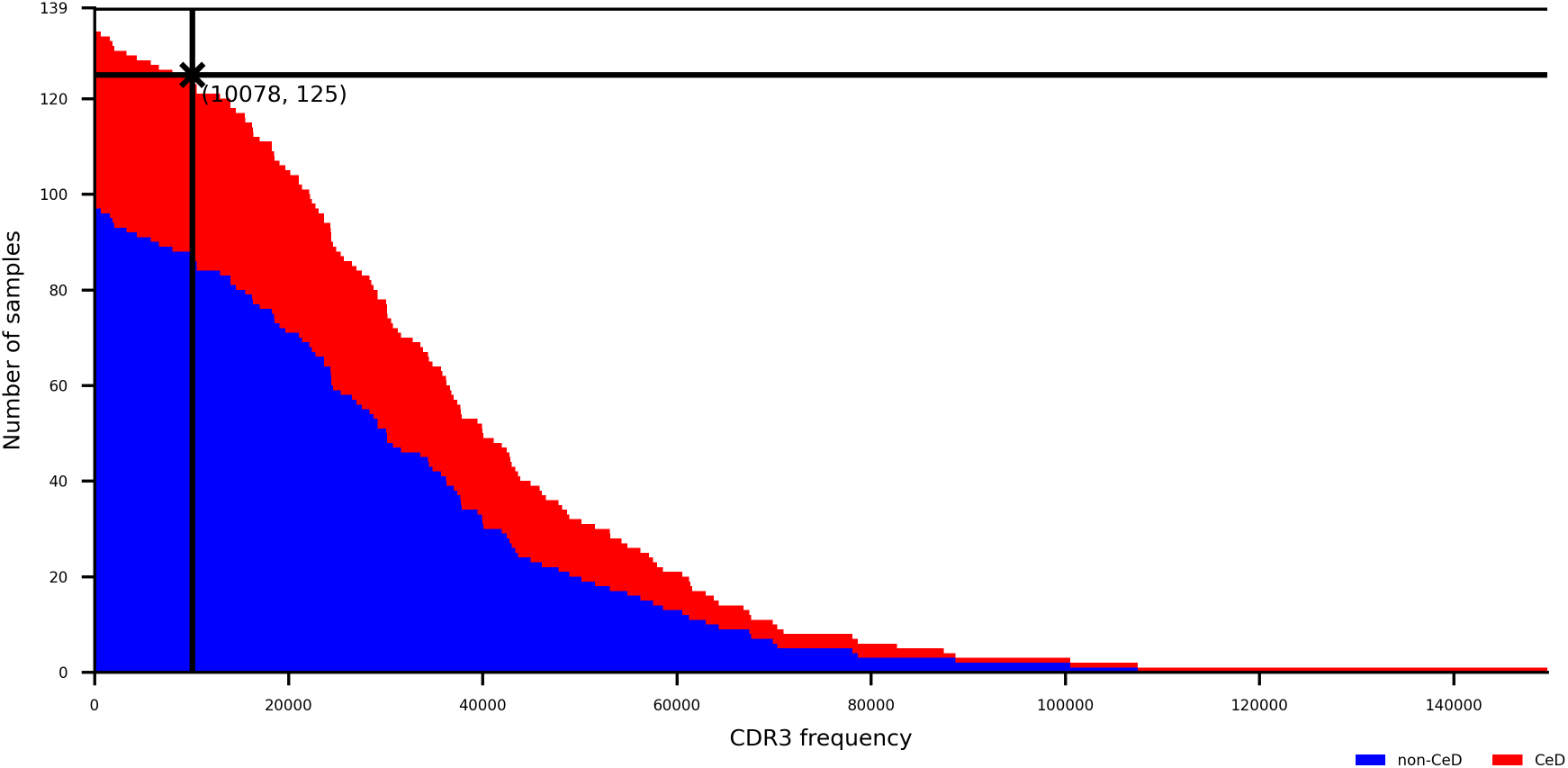
Step plot showing number of CeD training dataset samples that have at least a CDR3 count shown on horizontal axis, distinguished by class. A downsampling threshold is labelled with cross at 10078 sequences, retaining 125 samples.

**Table 4.** Summary of CeD datasets.

|  | CeD Training Dataset | CeD Testing Dataset |
| --- | --- | --- |
| Locus | TRD | TRD |
| Starting material | DNA from duodenal biopsies | DNA from duodenal biopsies |
| Samples before downsampling | 139 | 23 |
| CeD samples before downsampling | 38 | 11 |
| Non-CeD samples before downsampling | 101 | 12 |
| Downsampled reads to | 10078 | 10078 |
| Samples after downsampling | 125 | 22 |
| CeD samples after downsampling | 37 | 11 |
| Non-CeD samples after downsampling | 88 | 11 |

XGBoost models trained on 5 different kmer-based representations of the CeD training dataset are evaluated in validation, shown in table 5 and testing shown in table 6. Performance measures and the weight designated to each are consistent with those the the CMV classification problem, since class imbalance is also present in the CeD training dataset. In validation, BLOSUM62 reduced alphabet kmer features are estimated to result in better-performing XGBoost models across all measures, shown in table 5. Both kinds of kmer clusters result in AUROC over 0.8, and there is overlap of AUROC errors across features.

**Table 5.** Validation performance of XGBoost models on CeD testing dataset with five kmer representations.

| Features | AUROC | Accuracy | Sensitivity | Specificity | Balanced accuracy | MCC |
| --- | --- | --- | --- | --- | --- | --- |
| Kmers | 0.755 + -0.120 | 0.747 + -0.072 | 0.431 + -0.262 | 0.878 + -0.102 | 0.655 + -0.112 | 0.350 + -0.214 |
| Atchley reduced alphabet | 0.789 + -0.146 | 0.752 + -0.107 | 0.461 + -0.277 | 0.874 + -0.118 | 0.667 + -0.137 | 0.383 + -0.272 |
| BLOSUM62 reduced alphabet | **0.863** + -0.086 | **0.832** + -0.079 | **0.642** + -0.179 | **0.912** + -0.088 | **0.777** + -0.095 | **0.599** + -0.185 |
| Atchley kmer clusters | 0.811 + -0.074 | 0.749 + -0.070 | 0.514 + -0.225 | 0.848 + -0.075 | 0.681 + -0.105 | 0.376 + -0.198 |
| BLOSUM62 kmer clusters | 0.808 + -0.117 | 0.760 + -0.077 | 0.556 + -0.196 | 0.844 + -0.075 | 0.700 + -0.104 | 0.410 + -0.207 |

Each model is evaluated in 5-fold cross-validation repeated 3 times, and mean and standard deviation of performance measures including AUROC, accuracy, sensitivity, specificity, balanced accuracy and MCC for the testing folds are reported.

**Table 6.** Testing performance of XGBoost models on CeD testing dataset with five kmer representations.

| Features | AUROC | Accuracy | Sensitivity | Specificity | Balanced accuracy | MCC |
| --- | --- | --- | --- | --- | --- | --- |
| Kmers | **0.934** | **0.818** | 0.636 | **1.000** | **0.818** | **0.683** |
| Atchley reduced alphabet | 0.909 | **0.818** | **0.727** | 0.909 | **0.818** | 0.647 |
| BLOSUM62 reduced alphabet | 0.860 | **0.818** | 0.636 | **1.000** | **0.818** | **0.683** |
| Atchley kmer clusters | 0.760 | 0.682 | 0.545 | 0.818 | 0.682 | 0.378 |
| BLOSUM62 kmer clusters | 0.818 | **0.818** | **0.727** | 0.909 | **0.818** | 0.647 |

Each model is trained on entire CeD training dataset and tested on entire CeD testing dataset. Performance measures including AUROC, accuracy, sensitivity, specificity, balanced accuracy and MCC are reported for testing dataset.

In slight contrast to the validation performance, the standard kmer model results in the highest AUROC, balanced accuracy and MCC. The BLOSUM62 reduced alphabet model reaches the same balanced accuracy and MCC as the standard kmer model, but results in an AUROC of 0.860 which is lower than the best value of 0.934. The Atchley reduced alphabet model results in the same highest balanced accuracy of 0.818, and interestingly has a higher AUROC than its BLOSUM62 counterpart of 0.909, but a lower MCC. Both kmer cluster features result in lower AUROC than their reduced alphabet counterparts, though BLOSUM62 kmer clusters lead again to the highest balanced accuracy of 0.818.

Reduced alphabet sizes are set as a hyperparameter as for the CMV classification problem. Small reduced alphabets are chosen frequently when BLOSUM62 is specified as the basis of amino acid similarity, shown in fig 5. As shown in fig 5, a BLOSUM62 reduced alphabet of only size 6 is able to train a model that performs similarly to standard kmer features.

**Fig 5.**
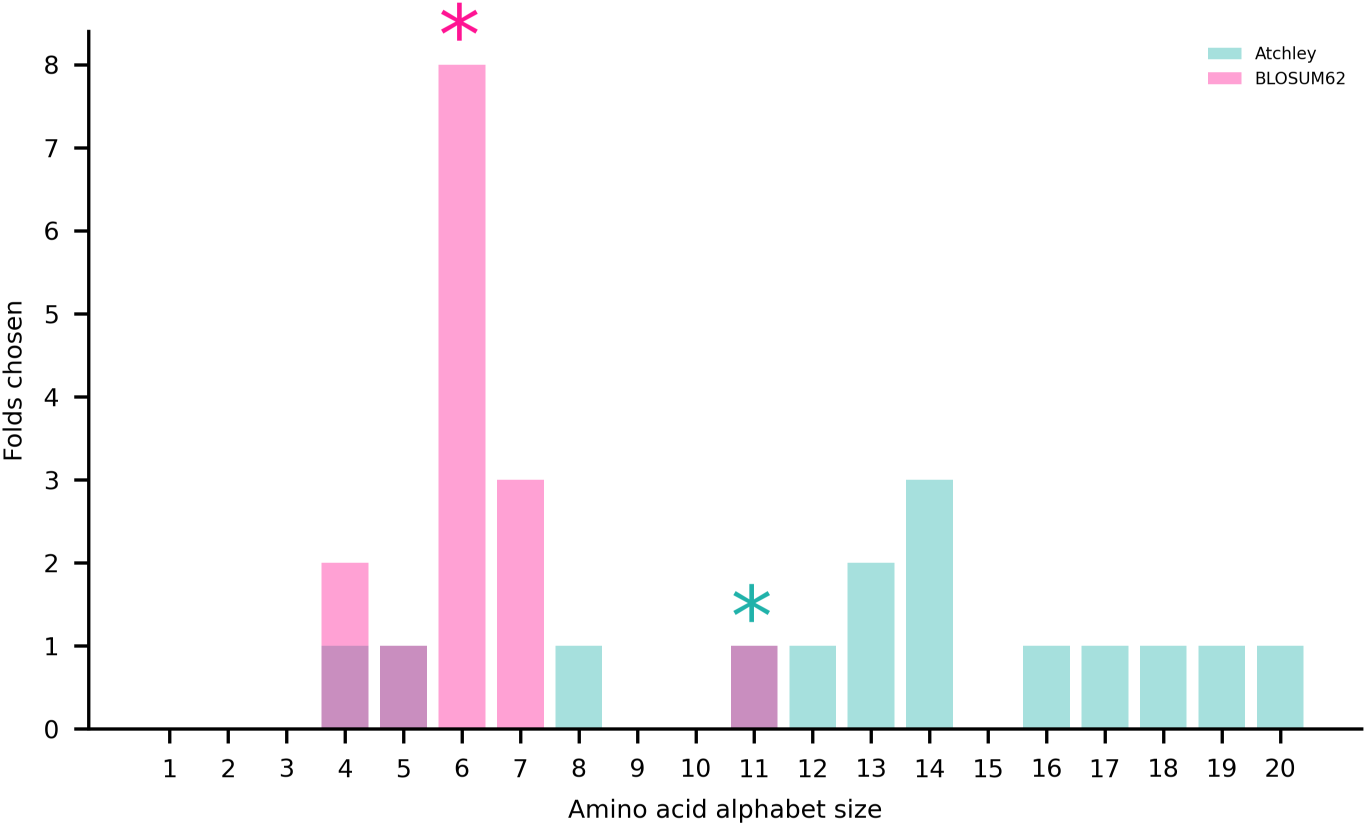
Reduced alphabet sizes chosen by XGBoost models in validation and testing with CeD datasets. Alphabet sizes chosen in training are indicated by asterisks.

XGBoost feature importance is obtained as for the CMV classification models. Fig 6 shows a summary of features used in the XGBoost model trained on the entire CeD training dataset. For standard kmer and reduced alphabet kmer features, over 100 features are incorporated into each model, an order of magnitude lower than the CMV XGBoost model. Many standard and reduced alphabet kmer features have a share of more than 1% of the total feature importance. Kmer cluster features tend to contain larger numbers of kmers, and fewer are utilised within a model. Within the annotated stacked bar plot showing feature importance as a proportion of the total in S2 Fig, the most important motifs within the reduced alphabet kmer models do not have obvious matches to the most important kmers within the standard kmer model. Also, the BLOSUM62 reduced alphabet model ranks motifs starting with [C] 2nd and 3rd, which may be starting motifs.

**Fig 6.**
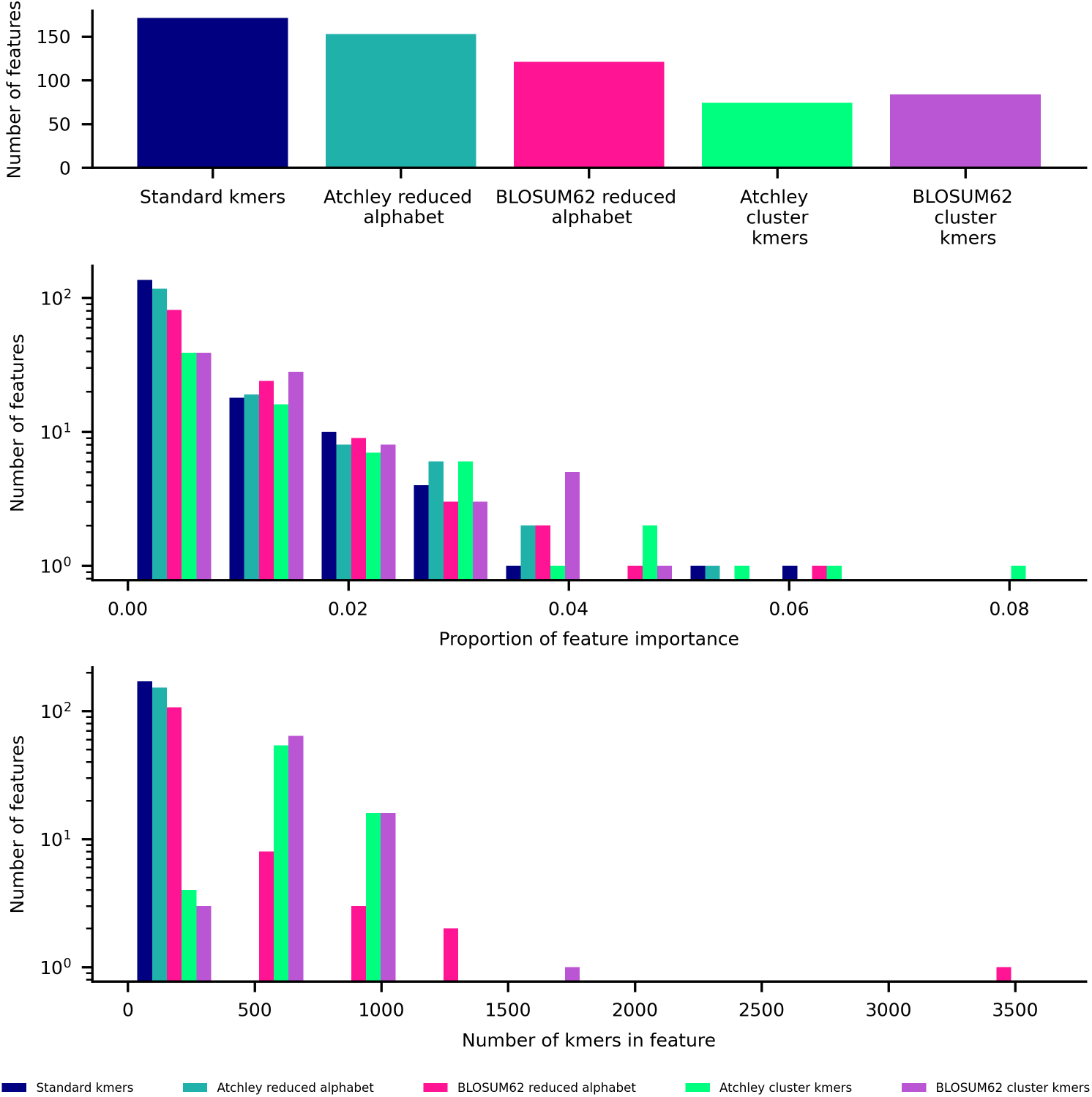
Summary of features used in XGBoost model trained on CeD training dataset for each feature type. Top: bar plot of number of features used in model. Middle: histogram of proportion of importance of features used in model. Lower: histogram of number of kmers within each feature used in model.

## Discussion

We introduce a novel and flexible kmer representation of the TCR reperoire based on a reduced alphabet of amino acids. To understand the utility of adding amino acid similarity information to a kmer representation of the TCR repertoire for classification, our approach is evaluated in comparison to the kmer cluster representation by Thomas et al. and standard kmers. Atchley factor and BLOSUM62 amino acid similarity specifications are assessed for both reduced alphabet and kmer clustering methods. The resulting 5 representations are evaluated in combination with XGBoost models for two different immune status classification problems: CMV infection and CeD status, which are each a means to provide evidence relating to immune status classification in general, as well as autoimmune disease classification.

For the CMV datasets, standard 4mer results very similar to those reported in the literature for 3mers are produced [7]. BLOSUM62 and Atchley factor reduced alphabet kmers may result in CMV status classification models that perform slightly better than standard kmers based on AUROC in testing. CeD classification models generally perform well as might be expected given that CeD datasets originate from intestinal tissue for which TCR repertoire differences between classes have previously been reported [41]. In validation on the CeD training set, a kmer representation that considers amino acid information seems to be beneficial, though testing indicates that a standard kmer model ultimately generalises better to the CeD testing set in terms of AUROC. Testing performance for CeD is greater than validation performance in most cases, which could be due to chance given its small size.

Kmer cluster approaches do perform reasonably well across classification models when 100 clusters of 4mers are used to represent the TCR repertoire, which would be equivalent to an alphabet size of roughly 3. However, it is not feasible to set the amino acid similarity threshold via number of clusters based on the training data when a full range of similarity thresholds are considered. The kmer clusters presented here are able to train models comparable to those by other kmer representations in some cases, but the ability to set amino acid similarity using our novel reduced alphabet approach has enabled insight to be gained about amino acid equivalences that result in the most effective classification basis for each combination of TCR locus and immune status that is investigated.

The observation that utilising amino acid alphabets of size 10 or smaller leads to just as good or a better representation for discriminating TCR repertoires by immune status than the 20 amino acids may be useful in understanding the utility of kmer representations in general. It may be that since a reduced alphabet reduces the dimensionality of the kmer representation, its effect is that of regularisation which would lead to improve generalisability. Alternatively, small reduced alphabets may allow flexibility in certain kmer positions, since one of many amino acids could be present. This flexibility may resemble a gap character, in which any amino acid could be present. The coeliac disease-associated R-motif observed in *αβ* TCR sequences contains gap characters [52], suggesting that a reduced alphabet approach may be a suitable tool for discovering other autoimmune disease-associated motifs.

The reduced alphabets of amino acids give rise to kmer-based features that can be described with a motif with known underlying amino acid relationships. In comparison to kmer clusters, where clusters are defined by a cluster centre which is a vector of attributes for k amino acids in the kmer as well as its cluster boundary, we reason that our reduced alphabet approach is more readily interpretable. Reduced alphabets lead to different classification bases than standard kmers, as in, divergent kmers or motifs are utilised to make predictions, which could lead to alternative biological interpretations. However, the feature importance results obtained from the XGBoost should be viewed with caution, since 5 different measures of importance: gain, total gain, cover, total cover and weight, could each be used to calculate feature importance, and these measures are inconsistent with each other [53]. Our results also highlight the challenge of interpreting XGBoost models trained on kmers whether a reduced alphabet is applied or not. 1000s of kmers or kmer-derived features may potentially be utilised in a model as is the case with the CMV training set, and interactions between these features are not explained by built-in feature importance measures.

For CMV status classification, representations utilising BLOSUM62 lead to higher AUROC than for Atchley factor representations. However, this is not always true for CeD status models. The CMV datasets, being larger in size, may provide a higher quality of evidence for the most appropriate amino acid similarity specification for TCR sequences. It could also be the case that the most appropriate amino acid similarity specification for *β* chain TCR sequences differs from *δ* chain sequences, since, in contrast to *αβ* T cells, *γδ* T cells are known to bind non-peptide antigens [54]. Further, properties or probable substitutions of amino acids that underlie function of TCRs may differ from those for proteins in general, and are a potential area of further study. Any amino acid relationships that are discovered as important in TCR binding with peptide-MHC complexes or antigens in general would be feasible to explore thoroughly in combination with a TCR repertoire classification model using our flexible reduced alphabet approach.

In addition, *δ* TCR chains from *γδ* T cells have longer CDR3 regions than those in TCR *β* chains of *αβ* T cells [55], so inclusion of interaction terms that combine 4mers or 4mer motifs may allow greater capacity to characterise TCR specificity within a classification model. However, interactions between features in XGBoost models cannot be understood using built-in feature importance measures, so it is difficult to ascertain whether a model essentially assembles longer discriminative motifs. In the future, kmer length could be fit as a hyperparameter alongside alphabet size to understand whether this is the case, which may lead to a model that is easier to interpret.

Overall, our work reveals important considerations when utilising an amino acid-aware kmer-based representation of the TCR repertoire in an immune status classification model. These results highlight the value of exploring TCR repertoire representations in a systematic manner, and provide a fast and flexible method for this exploration. Based on results for the CMV classification problem, we suggest that a small benefit may be gained in certain cases when employing a reduced alphabet representation over both a standard kmer representation and one based on kmer clusters. In addition, results indicate that greatly reduced alphabets of amino acids can provide as useful of a classification basis as the full amino acid alphabet. The ability to set amino acid similarity as a hyperparameter with up to only 20 discrete values enables amino acid relationships within kmers to be explored for different properties and substitution matrices in a way that would be challenging or unfeasible with a kmer clustering approach. However, the utility of TCR repertoire classification methods as a means to gain understanding of disease will ultimately only be realised if interpretability is priotitised in future work.

## Materials and methods

### TCR repertoire datasets

4 TCR repertoire datasets are used to evaluate models in this work, including training and testing datasets for each of CMV infection status and CeD status classification problems.

### CMV datasets

Two cohorts of TCR repertoire samples published in [4] were downloaded from the ImmuneACCESS database. For both cohorts, TCR *β* repertoires were obtained from genomic DNA in peripheral blood. Multiplex PCR was used for amplification, and sequencing was achieved using Illumina HiSeq. TCR repertoire samples were preprocessed by the original authors.

Cohort 1 was used as the CMV training dataset, and is stated to include 666 healthy bone marrow donors [4]. 665 samples were available for download, and 640 of these samples had CMV status labels. Two different column headings were identified as candidates from which to extract counts of each unique TCR sequence read, named *seq*_*reads* and *templates*. For 465 samples, only the *templates* column was present. For 123 samples, both *seq*_*reads* and *templates* headings were present, and both were populated with data. For another 76 samples, both headings existed but *templates* was not populated. A final single sample had both headings but missing values in the *templates* column. A decision to follow a procedure consistent with that by Katayama and Kobayashi [7], taking the *seq*_*reads* column if it exists and the *templates* column otherwise, was made for purposes of comparison.

Cohort 2 was used as the CeD testing dataset and includes healthy volunteers recruited to study infection [4]. 119 samples were available for download, which all included a templates columns rather than a *seq*_*reads* column.

### CeD datasets

The CeD training dataset includes TCR *δ* chain sequences from fully anonymised duodenal samples diagnosed with either active coeliac disease or diagnosed as normal (Ethics Ref: 04/Q1604/21, IRAS reference: 162057). DNA was extracted from the formalin-fixed paraffin embedded (FFPE) biopsy samples and bulk TCR *δ* chain libraries were created using the TCRD Gene clonality master mix (Invivoscribe) and then the Truseq Kit (Illumina). The final pooled library was sequenced by Illumina Miseq with 300 cycles. Paired end reads were preprocessed using the MiXCR align command [56].

Another *δ* chain dataset with coeliac disease status labels was previously obtained using similar methodology [28] and used as a CeD testing dataset. TCR *δ* chain sequences were obtained with DNA from FFPE duodenal biopsies. Despite some differences in preprocessing operations compared with the CeD training dataset, it is believed that this dataset can serve as a testing set due to similarity of preparation and sequencing.

In support of this publication, both CeD training and testing datasets are available at ImmPort (https://www.immport.org) under Study Accession SDY2976.

### Representing the TCR repertoire with kmer features

TCR repertoire datasets are represented with 5 different kmer definitions. Unaltered kmers, referred to as standard kmers throughout and described below, are evaluated as a null representation by which to compare representations that incorporate amino acid similarity information. Two different specifications of amino acid information are added to this kmer representation using both a reduced alphabet and kmer clusters as described by Thomas et al. [46]. Code used to generate these TCR repertoire representations, and subsequently to perform classification, are available at https://github.com/hannrko/enc_kmer_tcr_models.

### Kmers

A kmer or k-mer describes a short section of a sequence with length k, where kmers of a sequence overlap. In this work, kmers that are used to represent the TCR are made up of 4 amino acids, and each 4mer overlaps the previous 4mer by all but one amino acid. Evidence suggests that for *αβ* T cells, CDR3 positions in contact with antigenic peptides include 4 amino acids on average [43]. The choice of k as 4 throughout this work should therefore be appropriate for *β* chain CMV datasets. For *γδ* T cells, 4mers have been shown to separate samples in the CeD testing dataset by CeD status optimally [28], and so it is expected that 4mers should also be an appropriate representation for both CeD datasets.

### Amino acid encodings

Atchley et al. reduced a set of 54 recorded amino acid properties to 5 properties derived via factor analysis [47], which are termed Atchley factors. Substitution matrices such as BLOcks SUbstitution Matrices (BLOSUM) represent the chance that occurrence of amino acids at the same position in a pair of sequences is the result of relatedness rather than randomness [51, 57]. To represent amino acid similarities within kmers, both Atchley factors and BLOSUM62 are selected due to their frequent application to TCR repertoire representations [10, 11, 15, 42–46, 58, 59].

### Kmers defined with a reduced alphabet

Reduced alphabets of amino acids have been used to represent proteins in classification problems [60], where groups of similar amino acids are considered equivalent and represented by the same character. In this work, kmers are defined using a reduced alphabet of amino acids with variable size. The reduced alphabet is based upon grouping of the 20 naturally-occurring amino acids which is achieved through hierarchical clustering. Starting from a distance matrix representing pairwise distances between all 20 amino acids hierarchical clustering solutions are calculated using an average linkage policy, using SciPy [61] (version 1.11.4) functions from scipy.cluster.hierarchy. Over a full range of distance thresholds, each possible clustering solution is recorded to produce multiple reduced alphabets. Amino acids in all kmers are replaced by a character that is representative of a cluster of amino acids.

Frequencies of equivalent kmers that result are combined additively within each TCR repertoire sample.

The choice of reduced alphabet size can be treated as a hyperparameter. Since all reduced alphabets are stored in memory, only a single initial instance of hierarchical clustering is required to explore a full range of amino acid similarity thresholds. Aside from its size, the reduced alphabet representation is independent of the kmers that are observed in a TCR repertoire, since it is defined only on the basis of the 20 amino acids we expect to observe. This is in contrast to the kmer clustering described below, for which a clustering solution is dependent on kmers that are present in a sample, and that must be calculated separately for each similarity threshold specified.

Pairwise distances between amino acids are obtained in order to calculate the hierarchical clustering solutions used in this reduced alphabet approach. An Atchley factor-based distance matrix is defined through pairwise Euclidean distance of amino acids encoded by Atchley factors with standard scaling applied. A subset of BLOSUM62 corresponding to the 20 naturally-occurring amino acids was converted to a distance matrix. First, all elements of the matrix were made positive by subtracting the minimum value. Next, the matrix was normalised to have diagonal values of 1, as would be expected for a similarity matrix. To achieve this, each element representing substitution score of one amino acid for another was divided by the squared product of the self-substitution score of each amino acid. Finally, a distance matrix with diagonal elements of zero was retrieved by subtracting each element from 1.

### Kmer clusters

Kmer clustering is based on the approach described by Thomas et al. [46]. Unique kmers observed within a TCR repertoire training dataset are encoded by a vector of amino acid properties or features. The kmers are clustered into a specified number of clusters using K-means clustering with scikit-learn function sklearn.cluster.KMeans (version 1.4.2) [62] with all defaults except *n*_*clusters*. Kmers belonging to the same cluster are combined into the same feature additively, so that a TCR repertoire can be represented by the total counts of all kmers that map to each cluster. Kmers not observed in the training data are assigned to a cluster using the predict method. The number of clusters could conceivably be set as a hyperparameter, but the computational complexity of repeated K-means clustering limits the range of kmer similarity that can feasibly be explored. In this work, the number of clusters is set at 100 since strong performance was achieved by clustering 4mers into 100 groups by Thomas et al. [46].

Kmer clustering, in contrast to a reduced alphabet approach, takes encoded kmers as input. Atchley factors may be used to directly encode kmers as a vector for application of the kmeans clustering algorithm. Atchley factors are standardised prior to encoding, where each of 5 factors are transformed to have zero mean and unit standard deviation. The scaling applied should enable each factor to be considered in similar proportions during clustering. BLOSUM62 is converted to an encoding in a similar approach to that by Zhang et al. [42], where multidimensional scaling is applied to the BLOSUM62-derived distance matrix. 5 features from the result of multidimensional scaling are utilised for consistency with the Atchley factor encoding. The values of each of the five features are used to encode each amino acid in a kmer as with Atchley factors.

### Classification methods

Katayama and Kobayashi demonstrate that an improvement can be gained by training non-linear model over a linear model on a TCR repertoires represented by kmers [7]. We also aim to understand the benefit of a non-linear model with inclusion of amino acid similarity information, and therefore, we use gradient-boosted trees similar to [7]. Gradient-boosted trees were implemented using XGBoost (version 2.0.3) classification method XGBClassifier [63]. Parameters are *max*_*depth* = 10, *learning*_*rate* = 0.1, *n*_*estimators* = 100, and scale pos weight of negative to positive class ratio as recommended in the documentation, plus all other defaults. These parameters are set to values similar to defaults of LightGBM used by Katayama and Kobayashi since hyperparameter optimisation was revealed to contribute little to overall performance [7]. However, default max depth is unlimited in LightGBM which could lead to overfitting, and so a value of 10 was chosen as a conservative compromise between 6, the XGBoost default, and the unlimited default of LightGBM. This fixed set of hyperparameters allowed optmisation of reduced alphabet size to be prioritised.

### Optimisation of reduced alphabet size

For kmers with a reduced alphabet, the amino acid alphabet size was set based on performance on the training data. The performance of a model trained with each alphabet size, from 1 to 20, was evaluated using 5-fold cross validation with random shuffling. Performance is assessed using AUROC, where the alphabet size corresponding to the maximal value was chosen. If multiple optima were found, the smallest alphabet was selected from these, since there is a prior belief that reducing the dimensionality of the kmer representation will lead to less overfitting and improved performance on samples not in the training dataset.

### Evaluation of TCR repertoire classification models

The five different kmer representations of the TCR repertoire are assessed using TCR repertoire datasets including a training and testing set with cytomegalovirus status labels, as well as a training and testing set with coeliac disease status labels. TCR repertoire classification models are assessed using testing on separate testing datasets and 5-fold cross-validation repeated 3 times on training datasets. Performance measures, including AUROC, accuracy, sensitivity, specificity, balanced accuracy, and Matthew’s correlation coefficient, are recorded for all test folds. When performance is reported for cross-validation, scores are combined across folds by calculating the mean and standard deviation. In each training and testing fold, steps were taken to ensure no leakage of data, as follows: standard scaling was applied to features of all data, with mean and variance calculated using only data in the training folds; the size of reduced alphabet used to define kmers is set only based on performance on the data in training folds; kmers used to define kmer clusters were limited to those observed at least once within the data in the training folds; and testing data was held out until model development was finalised. Plots used to visualise classification model results are generated using Matplotlib [64].

## Supporting information

**S1 Fig.**
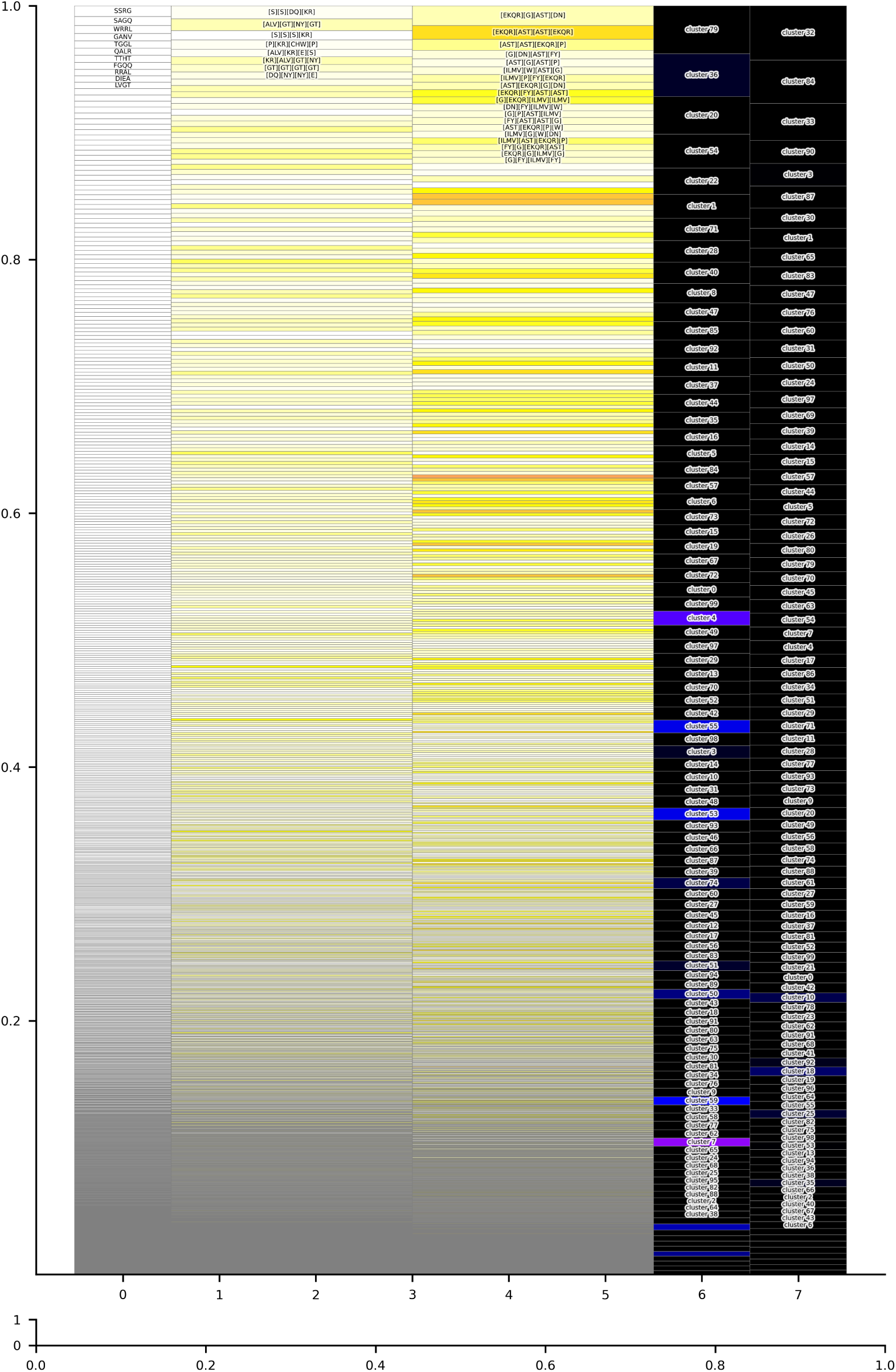
Importance of features of XGBoost model trained on CMV training dataset for each feature type. Feature importance, measured by gain, is shown in a stacked bar plot for each feature as a proportion of the total feature importance. Only features with at least 1% of total model importance are indicated by annotation. Background colour shows number of kmers included in each feature.

**S2 Fig.**
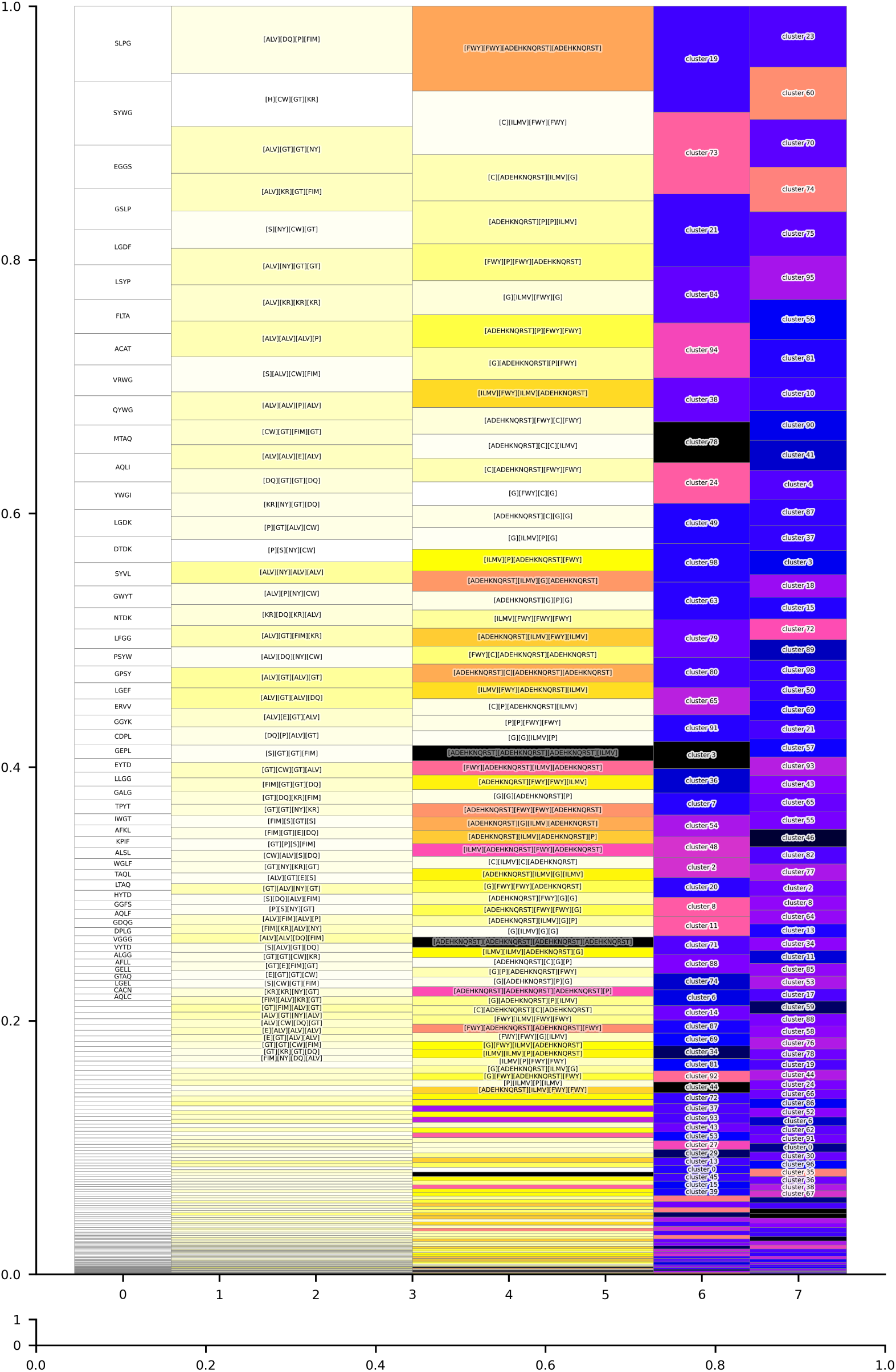
Importance of features of XGBoost model trained on CeD training dataset for each feature type. Feature importance, measured by gain, is shown in a stacked bar plot for each feature as a proportion of the total feature importance. Only features with at least 1% of total model importance are indicated by annotation. Background colour shows number of kmers included in each feature.

## Acknowledgments

This work was funded by Coeliac UK and Innovate UK (INOV01-18). H.K. received a Doctoral Training Studentship from EPSRC.

